# Unbiased counting of whole proteins with nanopores over the full dynamic range of a proteome: a computational study

**DOI:** 10.1101/2023.07.12.548698

**Authors:** G. Sampath

## Abstract

A computational model is presented for nanopore-based counting of intact single protein molecules in a sample regardless of the number of copies. It is based on measurable quantities and low detector bandwidths and can be applied to the full dynamic range of a proteome without the need for proteolysis or complex protein separation methods. Denatured unfolded whole protein molecules are assumed to translocate through a nanopore via electrophoresis and diffusion. A low solution pH helps keep the required detector bandwidth B in the 10-20 KHz range. An incremental Fokker-Planck drift-diffusion model is used to calculate two measurable quantities: 1) the total time (in the precision range set by B) for a protein to translocate through the pore, computed from the mean incremental translocation times of residues exiting the pore in succession; and 2) the volumes of protein segments inside the pore during translocation (used here as a proxy for the current blockade signal level) over alternate time blocks of width 1/2B. These are used to obtain volume-based string codes for each protein, substrings thereof as protein identifiers, and, if multiple copies are present, the copy number for a protein. This is a non-destructive single-molecule label-free alternative to mass spectrometry (MS) and other methods based on antibodies or optical tagging, it does not use any sequence identity information. Computational results are presented for the human proteome (Uniprot id UP000005640_9606; 20598 curated proteins). Total translocation times for the entire proteome (one copy per protein) are found to be in the tens of minutes. Over 80% of the proteome can be identified; higher percentages are possible by comparing whole proteins based on their string codes and total translocation times. Extrapolation of these results to a parallel 1000 pore array suggests that ∼10^9^ individual protein molecules can be counted in 15-20 hours.

## 1. Introduction

A major objective in next generation proteomics is obtaining the copy number distribution of proteins in single cells [1]. The range of copy numbers, commonly referred to as the dynamic range of a proteome, is spread over 7-10 orders of magnitude, with some proteins occurring in a single copy while others may have as many as 10^10^ copies per cell. Since proteins from both ends of this range have been implicated in disease conditions, it is desirable to isolate and identify, or better yet sequence, low abundance copy proteins. The present leader in protein sequencing and identification is mass spectrometry (MS), a bulk method that has in the past required hundreds of thousands to a million copies, although with progressive improvements in isolation and separation techniques, this number has continued to drop over the past decade [2]. Even as MS seeks to reduce sample sizes down to the attoliter level and below, there has been a growing interest in methods that can identify/count/sequence single protein molecules [3]. A variety of these methods have been developed over the last two decades, including nanopore-based and optical methods.

Sequencing may be done of whole protein sequences, of peptides obtained from the whole protein by proteolysis, from single residues cleaved from peptides, or from single residues cleaved from whole proteins. Protein sequencing is reviewed in [3,4]. Almost all of the methods mentioned therein are based on measuring one or more analog quantities. In a departure from this a binary-digital approach that can identify single terminal amino acids (AAs) successively cleaved from a peptide and bound with their cognate transfer RNAs has been proposed [5]. This method is virtually error-free as it is based on the extreme specificity (‘superspecificity’) with which tRNAs bind to their cognate AAs; it is yet to be translated into practice.

In contrast with protein sequencing, protein identification can often be done with a partial sequence, which is compared with a protein sequence database to determine if it occurs uniquely in some protein in the database [6-8]; this is usually the last step in a MS workflow. Other approaches have been reported in recent years. In [9] affinity-based reagents are used to identify target proteins; this is a bulk process. In [10] optical tags are attached to selected residues in short peptides; their fluorescence levels are measured and stored in a database for use in identification of an unknown peptide and its container protein. Protein identification based on peptides obtained by proteolysis is the mainstay of the Human Proteome Organization (HUPO) ‘s Human Proteome Project; the 2022 status report is at [11].

Sequencing/identification of whole proteins is preferred as it is non-destructive, it is also much more difficult than DNA sequencing for a number of reasons, not the least of which is that there are 20 AAs (as opposed to 4 bases in DNA), many of which have similar physical and chemical properties, making it hard to discriminate among them. Whole protein identification without proteolysis can be done in different ways. Some of these are based on: 1) The AA sequence; a partial sequence based on a few AAs is usually sufficient [12]; 2) Subsequence volumes [13]; 3) Antibodies or epitopes attached to the protein [9,14]; and 4) Measured sequences for the target sequence based on dividing the 20 AAs into 4 groups by AA volume: S(mall), M(edium), I(ntermediate), and L(arge) and measuring the blockade current level (which is a proxy for analyte exclusion volume) for successive residues in the protein sequence [15]. With 1, 2, and 4 it is assumed that successive individual residues can be distinguished (with varying degrees of difficulty) in the measured blockade current or other variable. At present measured translocation times for single residues are ∼1-10 μs, so this would require bandwidths of 100 Khz to 1 MHz or more.

Counting of individual proteins in a sample implies their identification. The centrality of protein counting in proteomics research is highlighted in [1,2]. Recent work in the area is briefly summarized below in Section 3.

### The present work

It is shown in theory that electrolytic cells (e-cells) with nanopores can be used to count every whole protein molecule in a sample regardless of the number of copies. A design-based approach is taken. Thus the counting procedure is based on analysis of the main parameters involved to determine what to compute and limit the main computation to measurable parameters. A simplified Fokker-Planck drift-diffusion model is used to calculate mean incremental translocation times for successive residues as they exit the pore. Translocation signals are divided into time blocks of length 1/2B, where B is the detector bandwidth. The volumes of the corresponding protein subsequences (which serve as a proxy for the current blockade signal) in alternate blocks are integrated, and the integrated volume sequences are used to extract protein identifiers for each protein. This is a non-destructive single-molecule label-free method that does not require complex protein separation and purification methods and is a potential alternative to mass spectrometry (MS), antibody-based identification, and optical methods based on fluorescent tags. In principle it can cover the full dynamic range of a proteome (∼10^10^ in single cells) in time spans comparable to those obtained with MS. Computational results for the human proteome (Uniprot id UP000005640_9606; 20598 curated proteins) show that with a transmembrane voltage of 50-100 mV and solution pH of 1 or 2, more than 80% of the proteins in the proteome can be identified. This can be further increased by analysis of pore sequence volume strings and sampled translocation signals of whole proteins. When extrapolated to a parallel 1000 pore array, these results suggest that 10^9^ individual protein molecules can be identified/counted in ∼15 hours.

## 2. The dynamic range problem and counting of whole proteins in a sample

Whether an expressed protein plays a significant role in the cell’s function (or malfunction) can be known only by looking at the full dynamic range of the cellular proteome. This range may be spread over 7-10 orders of magnitude, that is, some proteins may occur in as few as one copy, others may occur ten million times in the cell. In MS when an assay sample is subjected to analysis the protein molecules present in it are sampled and fed to a tandem LC-MS device [2]. Because of sampling bias, the more abundant proteins, which are usually present in larger numbers than low abundance proteins, are more likely to enter the spectrometer, while a protein that occurs in a small number of copies may be easily missed. Ongoing work on the sampling bias problem relies on sampling based on density variations in the purification and separation process that precedes a spectrometer so that the less abundant proteins have a better chance of entering the spectrometer [2].

In contrast with bulk methods like MS and gel electrophoresis, single molecule methods are based on inspecting single protein molecules in some detection space, which could be a nanopore in an e-cell or a glass slide to which the molecules carrying optical tags are attached. In principle nanopore-based counting should be able to solve the counting problem effectively because the pore acts as a sensor that examines a single molecule as it translocates through. Thus the protein molecules in the sample pass through the pore single file one after the other, so that in a perfect setting every molecule in the sample, regardless of the number of copies, would be counted.

As noted earlier, counting the number of copies of proteins in a sample presupposes knowing the identity of every protein in the sample. Several copy counting methods have been reported over the years, they are briefly summarized below.

In [16] nanopore-based single-molecule counting of dsDNA from SARS-Cov samples is discussed; DNA strands are identified by length and by comparison with sequences in a reference human genome database. In [17] target protein molecules in a complex sample are sampled by first attaching antibodies to protein molecules and conjugating the former with dsDNA strands. The antibody-DNA conjugates are released into solution and counted with a nanopore. In [18] label-based methods are described for controlled translocation of single protein molecules through nanopores aimed at counting and sequencing. Methods include oligonucleotide conjugation to protein molecules and use of molecular motors like ClpX. In [19] quantification of sample concentration through translocation rates (which are proportional to the number of molecules) is discussed. Such quantification usually requires knowledge of nanopore geometry and size or a calibration curve based on the latter. With the setup used it is shown that molecule translocation rate scales linearly with the baseline current for a fixed testing molecule and salt concentrations independent of pore geometry and voltage. An appropriate buffer is used to decrease the effect of electroosmotic flow. In [20] fluorescence spectroscopy is used to count proteins. Counting may be based on changes in intensity resulting from photobleaching or by comparing the measured fluorescent intensity with a standard reference protein. In [21] single molecule counting in solid-state nanopores is reviewed. In [22] microarrays are used to quantify proteins. In [1] the sampling of protein molecules from a single human cell (which contains about 6 × 10^9^ protein molecules) is reviewed. If the proteins are replaced with peptides obtained from proteolysis coverage of the full dynamic range would require reading ∼10^8^ peptides, and counting them would require identifying them individually. In [2] a comparative analysis of MS and single molecule methods for counting is presented. In particular the sampling bias issue is discussed at length. Thus LC-MS generates ion counts and uses them to estimate protein copy counts. LC-MS is at present able to count billions of peptide ions in a time span of 90 minutes. To avoid sampling bias the focus needs to shift to less abundant peptides. This can be done through advanced purification and separation methods. Thus high abundance proteins are thinned out, lower abundance proteins are separated, and the latter are entered separately into the spectrometer without being mixed in with the high abundance ones.

Image-based methods makes use of fluorescent tags attached to residues in the protein or peptide, with a different color tag used for different AAs. In [10] data capture proceeds in cycles, one per terminal residue, following by cleaving of the terminal residue with Edman degradation. Image-based methods have density limitations. Thus 10^6^ fluorophore tags would require an image area of 1 mm^2^, while 10^12^ would require an area of 1 m^2^ [2].

## 3. A nanopore-based counting model

Nanopore-based analysis of polymers is based on the use of an electrolytic cell with a thin membrane containing a nanodiameter pore that separates two chambers, commonly known as *cis* and *trans*, containing an electrolyte, typically KCl. When a voltage is applied across the membrane it ionizes the electrolyte and causes an electrolytic current due to K+ and Cl-ions traveling to opposite chambers (Figure 1). If an analyte (such as a polymer) molecule is introduced into the *cis* chamber, its monomers translocate through the pore to *trans* in sequence order, effectively causing a blockade of the normal or base electrolytic current. The blockade level can be considered a measure of analyte size, so that with sufficient precision it can be used to identify the analyte. If the analyte is a biopolymer like DNA, RNA, or protein/peptide, different monomers in the polymer may lead to different blockade levels, and the latter, ideally, may be used to sequence the polymer.

**Figure 1.**
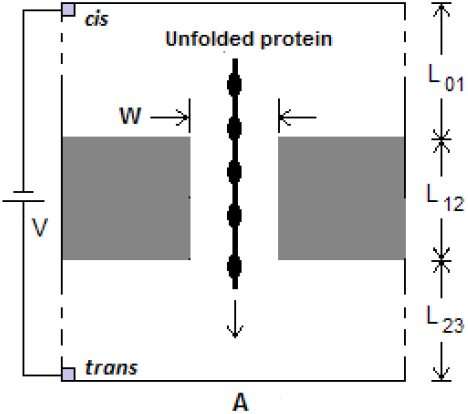
Schematic of electrolytic cell (e-cell); membrane containing nanopore separates *cis* and *trans* chambers containing salt (KCl) solution. Ionic current flow in e-cell is due to K^+^ and Cl^-^ ions; analyte translocating through pore causes reduction in open pore current; 98% of voltage V between *cis* and *trans* drops across pore. In present study length of pore L = 5-10 nm; voltage V is in 0-05-0.1 range; W = diameter of pore, typically 4-8 nm.

This section takes a design-based approach to protein counting with nanopores. There are two parts:

1. Parameter properties: A number of analyte parameters have to be taken into account in a nanopore-based counter. They include:
  i. the diffusion coefficient of a protein when it is in the constrained region of the pore;
  ii. electrical charge carried by residues and their mobility inside the pore, and relatedly solution pH; and
  iii. applied voltage.
2. Analytical properties: Only measurable properties are to be used in the analytical procedure used. These are:
  i. protein translocation times through the pore;
  ii. current blockade levels due to the segment of the protein inside the pore.

### Parameter analysis

#### 1. Diffusion coefficient

The diffusion coefficient of a molecule is given by Equation 1:

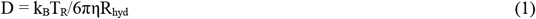

with k_B_ = Boltzmann constant (1.3806 × 10^−23^ J/K), T_R_ = room temperature (298° K), η = solvent viscosity (0.001 Pa.s for water), R_hyd_ = hydrodynamic radius of the analyte molecule (Å). Table 1 gives the R_hyd_ values for the 20 AAs:

**Table 1.** Hydrodynamic radii of the 20 AAs and their diffusion coefficients. R_aa_ values 10^−10^ m, D_aa_ values 10^−10^ m^2^/s

|  | A | R | N | D | C | Q | E | G | H | I | L | K | M | F | P | S | T | W | Y | V |
| --- | --- | --- | --- | --- | --- | --- | --- | --- | --- | --- | --- | --- | --- | --- | --- | --- | --- | --- | --- | --- |
| $R_{aa}$ | 2.66 | 3.6 | 2.98 | 3.02 | 2.86 | 3.23 | 3.14 | 2.32 | 3.49 | 3.24 | 3.39 | 3.69 | 3.08 | 3.35 | 2.68 | 2.76 | 3.04 | 3.5 | 3.57 | 3.32 |
| $D_{aa}$ | 8.21 | 6.06 | 7.32 | 7.23 | 7.63 | 6.76 | 6.95 | 9.41 | 6.25 | 6.74 | 6.44 | 5.91 | 7.09 | 6.52 | 8.14 | 7.91 | 7.18 | 6.24 | 6.11 | 6.57 |

In [23] DNA is considered a stiff rod with an axial diffusion coefficient D of about D = 68 ×10^−12^ m^2^/s. In [24] D for a confined polymer with N = 12 residues has an upper bound of 0.5-10 ×10^−12^ m^2^/s, which is at least 2 orders of magnitude smaller than the bulk diffusion coefficient for a single strand of DNA with the same number of bases. See [25] for a log-log plot of D versus polymer chain length N = 23, 31, 45, 57, and 73, the plot has a slope of -0.93.

For a rigid rod of length L and radius R, the hydrodynamic radius is given by

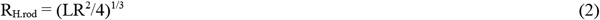

In a recent study [26] the experimentally obtained D value for a peptide with 10 residues is given as (0.6-1.2) × 10^−12^ m^2^/s inside the biological pore alpha-Hemolysin (APH). In the present study with a pore length of 5 nm and a profile length (= dimension of residue along the polymer chain) of 5 Å for every residue, a D value of 0.5 ×10^−12^ m^2^/s is considered.

#### 2. Mobility and pH

Consider the set of 20 standard AAs:

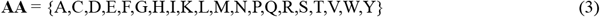

Every isolated AA is charged because of the charge in the amino (N-) and carboxyl (COOH-) terminals [27]. Only five AAs, namely C, D, E, H, K, R, and Y, are intrinsically charged because of the electrical charge in their side chains. The charge carried by an AA is equal to C_mult_q, where 0 ≤ |C_mult_| ≤ 2 and q is the (absolute) electron charge. C_mult_ depends on the pH level of the solution. In the present study the five naturally charged AAs are of significance. The other 13 do not play a role when they are internal to the polymer chain. Table 1 shows the dependence of electrical charge on pH for the charged AAs for selected pH values.

The mobility of a molecule in an electric field is given by Equation 4:

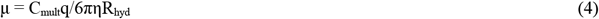

where q = electron charge (−1.619 × 10^−19^ coulomb), and C_mult_ = charge multiplier for the analyte as discussed above.

Table 2 gives the mobility values of the charged AAs at the pH values in Table 1.

**Table 2.** C_mult_ values of the charged AAs at selected pH values (Amino acids not listed have C_mult_ = 0.)

| pH | R | D | C | E | H | K | Y |
| --- | --- | --- | --- | --- | --- | --- | --- |
| 0 | 1.0000 | -0.0002 | 0.0000 | -0.0001 | 1.0000 | 1.0000 | 0.0000 |
| 1 | 1.0000 | -0.0022 | 0.0000 | -0.0006 | 1.0000 | 1.0000 | 0.0000 |
| 2 | 1.0000 | -0.0219 | 0.0000 | -0.0056 | 0.9999 | 1.0000 | 0.0000 |
| 7 | 1.0000 | -0.9996 | -0.0620 | -0.9982 | 0.0909 | 0.9997 | -0.0009 |
| 12 | 0.7512 | -1.0000 | -0.9998 | -1.0000 | 0.0000 | 0.0328 | -0.9884 |
| 13 | 0.2319 | -1.0000 | -1.0000 | -1.0000 | 0.0000 | 0.0034 | -0.9988 |
| 14 | 0.0293 | -1.0000 | -1.0000 | -1.0000 | 0.0000 | 0.0003 | -0.9999 |

**Table 3.**
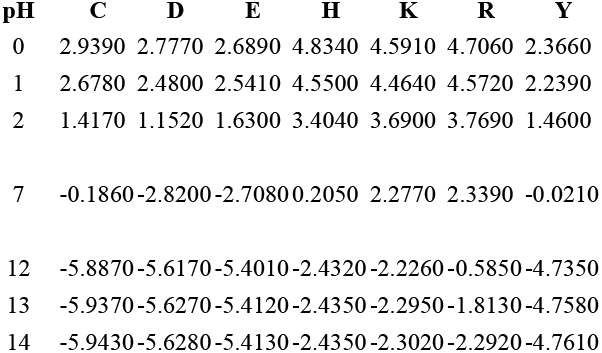
μ (mobility) values of the charged AAs at selected pH values (10^−8^ m^2^/s) (Amino acids not listed have μ = 0.)

| pH | C | D | E | H | K | R | Y |
| --- | --- | --- | --- | --- | --- | --- | --- |
| 0 | 2.9390 | 2.7770 | 2.6890 | 4.8340 | 4.5910 | 4.7060 | 2.3660 |
| 1 | 2.6780 | 2.4800 | 2.5410 | 4.5500 | 4.4640 | 4.5720 | 2.2390 |
| 2 | 1.4170 | 1.1520 | 1.6300 | 3.4040 | 3.6900 | 3.7690 | 1.4600 |
| 7 | -0.1860 | -2.8200 | -2.7080 | 0.2050 | 2.2770 | 2.3390 | -0.0210 |
| 12 | -5.8870 | -5.6170 | -5.4010 | -2.4320 | -2.2260 | -0.5850 | -4.7350 |
| 13 | -5.9370 | -5.6270 | -5.4120 | -2.4350 | -2.2950 | -1.8130 | -4.7580 |
| 14 | -5.9430 | -5.6280 | -5.4130 | -2.4350 | -2.3020 | -2.2920 | -4.7610 |

Translocation of a protein through the pore is determined in part by the mobility of the subsequence occupying the pore because 98% of the applied voltage drops across the pore. Subsequence mobility is determined by the total charge contained in the segment of the protein inside the pore. Consider a protein P with N residues given by

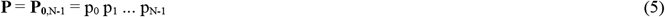

A segment with M residues inside the pore given by **P** _k,k+M-1_ = p_k_ p_k+1_ … p_k+M-1_ has a total charge and mobility given respectively by

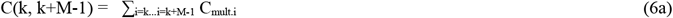

and

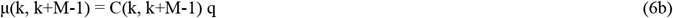

where C_mult.i_ is the charge multiplier for the k-th residue in the segment. Depending on the residues making up the subsequence C(k, k+M-1) can be anywhere between 0 and M (see Table 2; all AAs not in the table have C_mult_ = 0).

#### 3. Residue and subsequence volumes

Blockade levels are roughly proportional to the volume of the subsequence occupying the pore. The latter keeps changing as the residue at the *trans* end of the pore shifts out into *trans* and the next residue after the *cis* end enters from *cis*. Residue volumes are given in Table 4, they are taken from [28].

**Table 4.** Mean volumes of the 20 standard AAs (10^−9^ m^3^)

| AA | G | A | S | C | D | T | N | P | V | E |
| --- | --- | --- | --- | --- | --- | --- | --- | --- | --- | --- |
| Mean vol | 59.9 | 87.8 | 91.7 | 105.4 | 115.4 | 118.3 | 120.1 | 123.3 | 138.8 | 140.9 |
| AA | Q | H | M | I | L | K | R | F | Y | W |
| Mean vol | 145.1 | 156.3 | 165.2 | 166.1 | 168 | 172.7 | 188.2 | 189.7 | 191.2 | 227.9 |

### Analytical properties

For a nanopore-based counting method to succeed across the full dynamic range of a proteome the following set of conditions has to be met:

1. The rate at which a whole protein translocates through the pore must be compatible with the bandwidth of the detector. Within this band there should be no missed molecules, except perhaps for a small subset of outliers. This may require slowing down the analyte as it translocates through the pore. This problem is addressed in detail in [29].
2. The blockade signals obtained must have sufficient information that can be used to identify all copies of a distinct protein as being the same protein, and to distinguish it from all the other proteins. The method described here appears to meet both the above conditions, with a small number of microproteins (proteins with 20 or fewer residues) excluded from its purview.
3. Protein identification can only be based on measurements made within the given bandwidth B.

#### 1) Translocation time

An F-P equation is used to model analyte motion in the e-cell in Figure 1A and compute the dwell/translocation time of an analyte in/through the pore. Let T be the time of translocation of an analyte through the pore and E(T) its mean. Let D be the diffusion constant, μ the analyte mobility, and v_z_ = μV_12_/L the drift velocity through the pore. With α = v_z_L/D = μV_12_/D, V_12_ = V, and L_12_ = L, it can be shown that

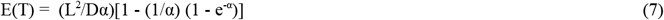

The following three cases can be considered:

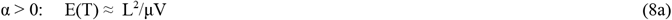

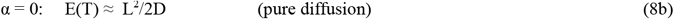

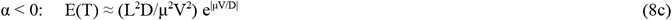

For the mathematical analysis leading to Equations 7 and 8 see the Supplement to [30].

Now consider a whole protein translocating from *cis* to *trans* through the pore. Neglecting end effects (the initial time to taken to fill the pore on entry from *cis* and the final time to empty the pore as the protein exits fully into *trans*, the time taken for the segment inside the pore **P** _k,k+M-1_ = p_k_ p_k+1_ … p_k+M-1_, k=0,1,…N-M+1 to translocate out can be considered in terms of the times taken by each residue as it exits the pore. This time can be written as

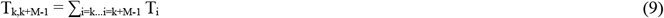

where T_i_ = time for residue i at the *trans* end of the pore to move from inside the pore into *trans*. T_i_ can be approximated from the time taken for the segment **P** _k,k+M-1_ to move out as a whole into *trans*. The latter is calculated from Equation 8 by assuming the segment to be a rigid rod that behaves like a particle with length L and mobility given by Equations 6 and 4. Which parts of Equations 7 and 8 apply depend on the mobilities of the residues p_k_, p_k+1_, … p_k+M-1_ inside the pore:

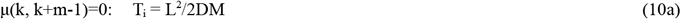

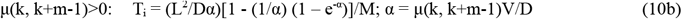

This incremental computation allows capturing the variations in translocation speed of the segment due to the changing amount of charge due to the residue at the end of the pore exiting and another entering the pore from *cis*.

Neglecting end effects the total translocation time for a whole protein **P** is given by

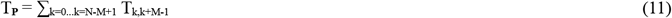

Figure 2 shows the blockade volume (see below) due to the protein segment inside the pore vs the translocation time of the residue exiting at the *trans* end of the pore. The top panel shows the raw translocation signal for protein number 5 (this is the the sixth protein, numbering starts with a base value of 0) in the human proteome UP000005640_9606, (protein id A6NL46, 340 residues). The bottom panel is what a detector with unlimited bandwidth and unlimited current precision would see.

**Figure 2.**
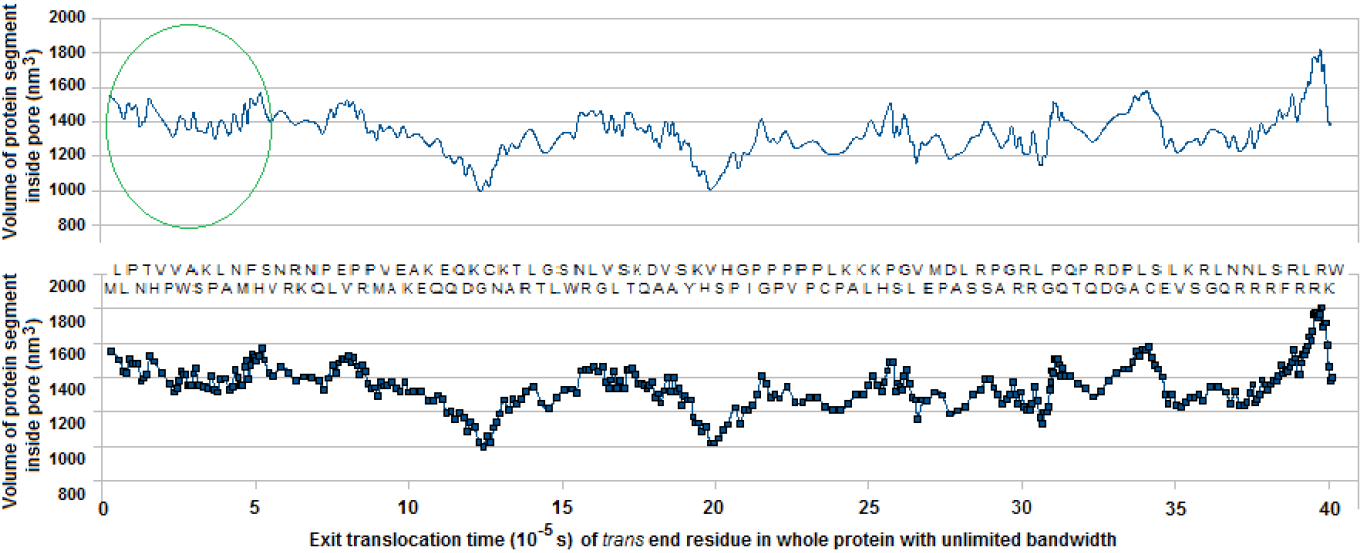
Blockade volume vs translocation time of *trans* end residue in pore of length 5 nm. Top panel shows raw signal. Bottom panel shows signal as seen by detector with unlimited bandwidth and unlimited current precision. Computed data are for protein number 5, id A6NL46, 340 residues, in human protein UP000005640_9606. Total translocation time for whole protein = 4.01 × 10^−4^ s

Translocation times and blockade volumes for protein 5 are given in Supplementary File 4.

#### 2) Subsequence volumes as a proxy for blockade levels

The current blockade caused by the translocating protein varies with the volume of the segment occupying the pore at any time. Similar to the calculation of translocation times through the pore in terms of incremental translocation times of successively exiting residues the volume of a segment occupying the pore can be calculated in terms of the change in volume due to successive residues exiting the pore. This leads to

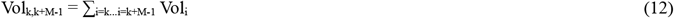

where Vol_i_ is the volume of residue i. Residue k is the next residue to exit into *trans*, so that in the next round k becomes k+1 and Vol_k_ is replaced with Vol_k+1_ and residue k+M enters the pore. This can be written as a difference equation:

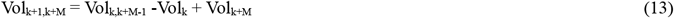

Each of the volumes Vol_i_ in Equation 9 contributes to the total volume over the time T_i_. This time is usually much smaller than 1/2B and cannot be sensed by the detector, which can only make measurements over a time period of width 1/2B or more; see Figure 3. The volume that is sensed by the detector during the q-th pulse [q/B, (2q+1)/2B],q=0,1,2… can be approximated with

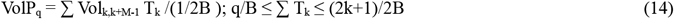

**Figure 3.**
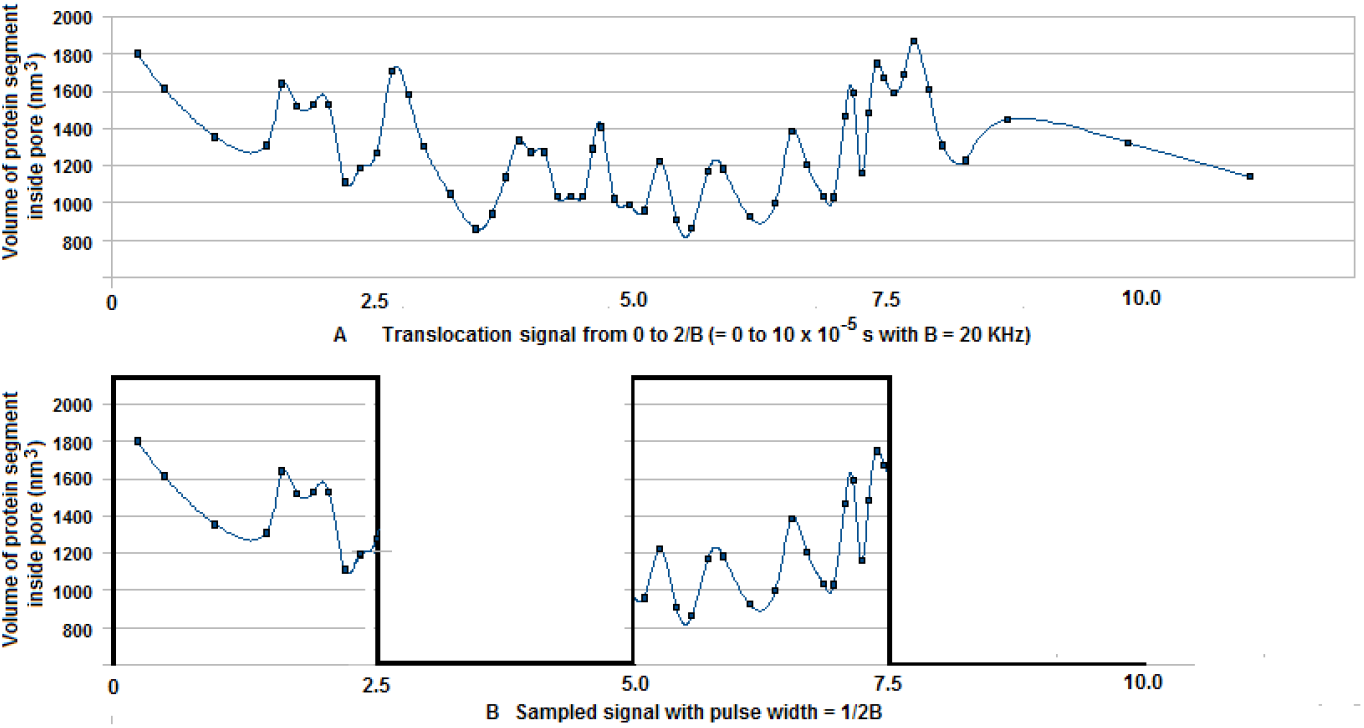
Sampling of translocation signal by band-limited detector with bandwidth B = 20 KHz.

**Figure 4.**
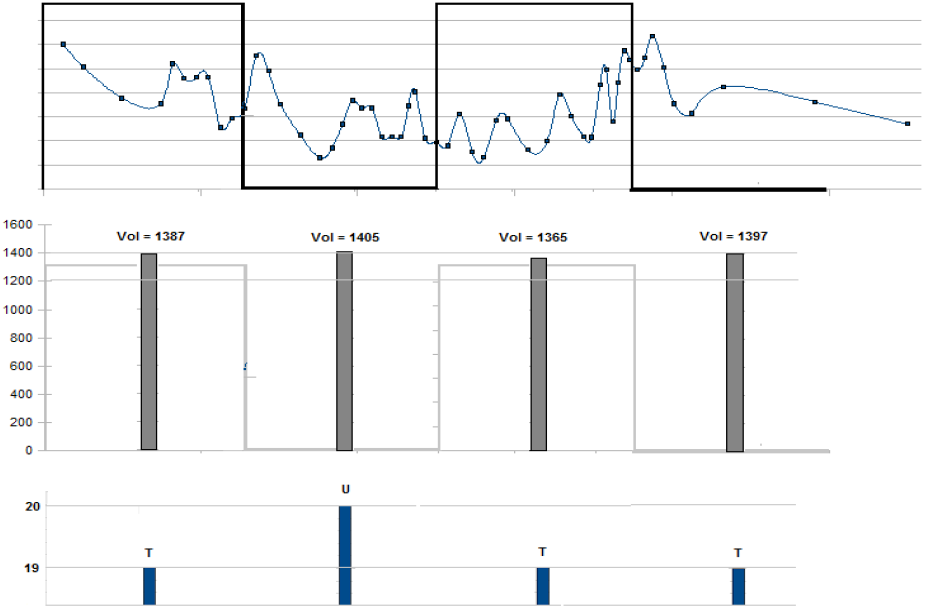
Digitization of sampled signal followed by translation to character string. Total volume in each half-period in top panel is shown in middle panel. Bottom panel shows translated characters for digitized volumes.

The top part of Figure 3 shows the calculated individual volumes of successive exiting residues and their times. The bottom part of the figure shows the sampling of this time sequence by the detector with a pulse width of 1/2B. The volume measured by the detector is the total volume over the time period 1/2B. (The next time period of 1/2B is skipped in the sampling process.) Thus the output of the detector is the sequence of values obtained by integrating over the incremental volumes during the pulse of width 1/2B. This output acts as the protein’s signature, from it one or more identifiers for the protein have to be extracted. This is discussed in the following paragraphs. (Notice that the number of residues whose volumes contribute to the total volume during a sample pulse may vary considerably. For example in Figure 3 it is 10 in the first pulse and 17 in the second. The sequence of integrals of the curve in Figure 3 between q/B and (2k+1)/2B for q = 0, 2, … can be used as the protein’s signature.

VolP_q_ can be digitized based on the volume resolution available. If analyte exclusion volumes are measured with a precision of Vol_step_ (nm^3^), the digitized volume of the subsequence inside the pore in the q-th pulse is then written as

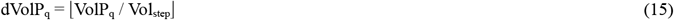

A Vol_step_ value of 70 nm^3^ is used here, it is based on [15].

The protein sequence is now represented by a digitized volume sequence:

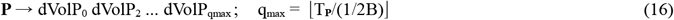

It is convenient to work with character strings so this sequence can be converted into an ASCII character string with the mapping

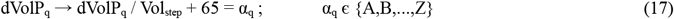

As an example consider protein 5, id A6NL46 in the human proteome (Supplementary File 1). The sequence has 340 residues:

MRLCLIPWNTTPHRVLPPVVWSAPSRKKPVLSARNSMMFGHLSPVRNPRLRGKFNLQLPSLDEQVIPTRLPKMEVRAEEPKEATEVKDQVETQGQEDNKTGPCSNGKAASTSRPLETQGNLTSSWYNPRPLEGNVHLKSLTEKNQTDKAQVHAVSFYSKGHGVTSSHSPAGGILPFGKPDPLPAVLPAPVPDCSLWPEKAALKVLGKDHLPSSPGLLMVGEDMQPKDPAALRSSRSSPPRAAGHRPRKRKLSGPPLQLQQTPPLQLRWDRDEGPPPAKLPCLSPEALLVGKASQREGRLQQGNMRKNVRVLSRTSKFRRLRQLLRRRKKRWQGRRGGSRL

Its volume coded character string has 8 characters: VURTRRSS

Volume code strings for all the proteins in the human proteins except for a small number of outliers are listed in Supplementary File 2.

#### 3) Protein identification

In [15] the set of AAs is divided into subsets based on AA volume and labeled S(mall), I(ntermediate), and L(arge) and the protein sequence relabeled with the code (S,I,M,L). Identifiers are then obtained from the relabeled sequence. This implicitly assumes that single residues in the protein sequence can be detected by the detector as the protein translocates through the pore. As noted earlier this would require bandwidths of 1 MHz or more.

In the present study protein identification is based on:

A. using substrings of the coded string in Equations 16 and 17; if this substring is not present in the coded strings of all other proteins it is an identifier for this protein;
B. comparing the full coded string for a protein with those of the other proteins.

Methods (A) and (B) are considered next. In either case it is more efficient to compare proteins with similar properties rather than to compare a protein with all the other proteins regardless of wide differences in these properties. Such properties include: 1) length of protein as given by the number of residues; and 2) translocation times of whole proteins through the pore.

Figure 5 shows the frequency distribution of proteins in UP000005640_9606 by number of residues.

**Figure 5.**
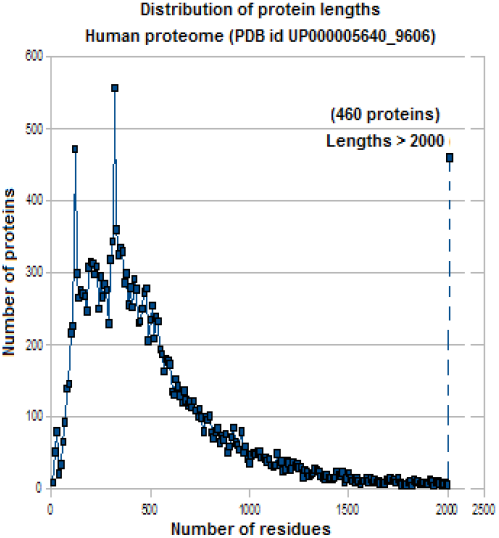
Length (no. of residues) distribution of proteins in human proteome (Uniprot id UP000005640_9606; total 20598 proteins)

However, the number of residues is not a measurable quantity, so it will not be considered further. Translocation times on the other hand are measurable. Figure 6 shows the total translocation time, maximum translocation time, and minimum translocation time for the proteins of UP000005640_9606 for different values of the transmembrane voltage V and solution pH.

**Figure 6.**
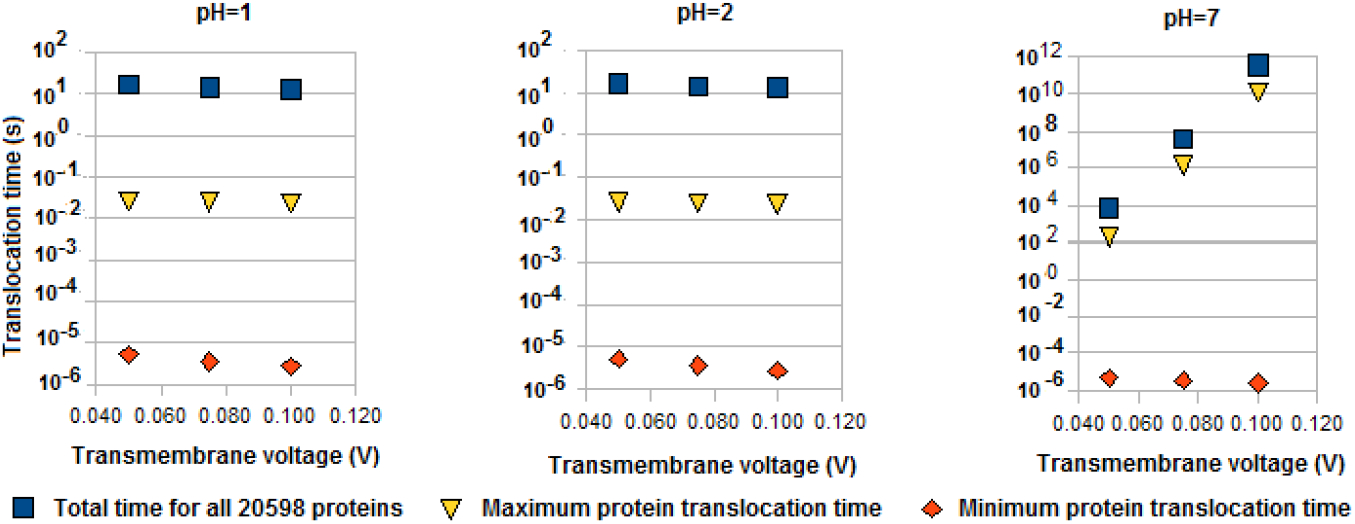
Translocation times for proteins in human proteome for three different transmembrane voltages and three values of solution pH

Notice how with normal pH (physiological pH = 7) the translocation times can be extremely large, they are in the 10^2^ – 10^12^ s range. This is caused by the D and E residues, which have an electrical charge that is opposite to the transmembrane electric field and causes the protein to be delayed by very large amounts (which arise from large values of μV/D in Equation 8c, here D refers to the diffusion coefficient) and may even cause its regression into *cis*. With a low pH value of 1, the C_mult_ value for D (the AA) is -0.0022 (see Table 2), compared with -0.9996 at pH = 7. (For the reason why a low pH value is chosen rather then a high one, although qualitatively the latter has the same effect as the former, see Item 5 in the discussion in Section 4.)

##### (A) Partial sequences as protein identifiers

Equation 17 leads to the coded volume string

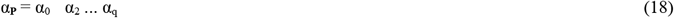

where the α’s are characters from the alphabet A-Z.

Substrings of α_**Pi**_ for one protein can now be compared with those for another protein α_**Pj**_ using string search algorithms. Comparisons may be unconstrained, with a substring for a protein’s codestring compared with the substrings of the code string of every other protein. Or it may be constrained to proteins that have similar characteristics. For example it does not makes sense to look for substrings from a short code string to one in a long code string, as the two proteins are obviously different even if they contain the same substring(s). The length of the code string, which is roughly proportional to the translocation time of the whole protein, can be used to determine whether to do the match or not. Figure 7 shows identification rates with unconstrained matching and matching of proteins that are within 1/B of the translocation time of each other.

**Figure 7.**
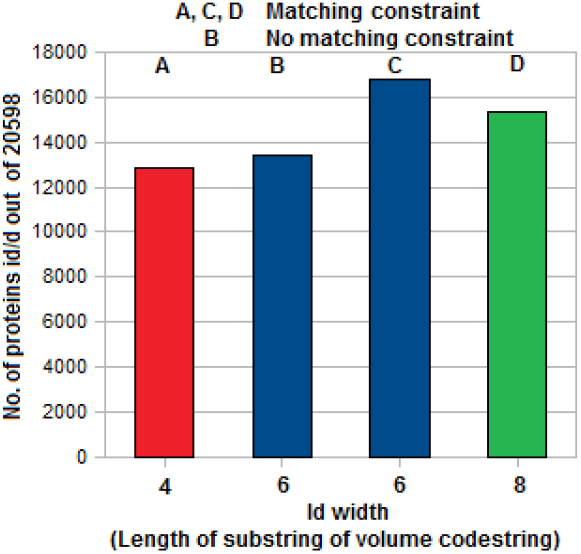
Distribution of number of protein ids in volume code string for every protein in the human proteome for different id lengths in code string. Ids obtained by matching substrings of length 4-8 against substrings of other proteins in proteome. In A, C, and D only proteins with code string lengths within 1/B of each other are compared. In B there is no constraint, substrings of a protein of given id length are compared with substrings of the same length in every other protein.

Ids of length 6 with a nanopore of length 5 nm, transmembrane voltage of 0.05 V and pH = 1 for all proteins in the human proteome except for a few outliers (such as microproteins with 20 or fewer residues) are listed in Supplementary File 3.

###### Phase difference in a measured translocation signal

An issue to be considered in practice is the exact time when sampling starts. Synchronizing this with the first residue of the protein when it enters the pore is not possible in practice. Viewing the sampling start time as an offset or phase from the first residue, subsequence volumes can be considered for different offsets and volume code strings obtained for each phase for up to k residues. Column 2 in Table 6 shows the code strings for protein 5 for phase differences k = 0,1,…, 19. The code strings for the 20 phases are seen to be fairly similar. Matching of a substring of the code string for a given protein now needs to be done with substrings in the code strings for all 20 phases.

One way to estimate the phase is to attach leader and trailer peptides to the protein. Consider protein 5:

MRLCLIPWNTTPHRVLPPVVWSAPSRKKPVLSARNSMMFGHLSPVRNPRLRGKFNLQLPSLDEQVIPTRLPKMEVRAEEPKEATEVKDQVETQGQEDNKTGPCSNGKAASTSRPLETQGNLTSSWYNPRPLEGNVHLKSLTEKNQTDKAQVHAVSFYSKGHGVTSSHSPAGGILPFGKPDPLPAVLPAPVPDCSLWPEKAALKVLGKDHLPSSPGLLMVGEDMQPKDPAALRSSRSSPPRAAGHRPRKRKLSGPPLQLQQTPPLQLRWDRDEGPPPAKLPCLSPEALLVGKASQREGRLQQGNMRKNVRVLSRTSKFRRLRQLLRRRKKRWQGRRGGSRL

With a header of 100 E’s and a trailer of 50 E’s the resulting sequence is

EEEEEEEEEEEEEEEEEEEEEEEEEEEEEEEEEEEEEEEEEEEEEEEEEEEEEEEEEEEEEEEEEEEEEEEEEEEEEEEEEEEEEEEEEEEEEEEEEEEEMRLCLIPWNTTPHRVLPPVVWSAPSRKKPVLSARNSMMFGHLSPVRNPRLRGKFNLQLPSLDEQVIPTRLPKMEVRAEEPKEATEVKDQVETQGQEDNKTGPCSNGKAASTSRPLETQGNLTSSWYNPRPLEGNVHLKSLTEKNQTDKAQVHAVSFYSKGHGVTSSHSPAGGILPFGKPDPLPAVLPAPVPDCSLWPEKAALKVLGKDHLPSSPGLLMVGEDMQPKDPAALRSSRSSPPRAAGHRPRKR KLSGPPLQLQQTPPLQLRWDRDEGPPPAKLPCLSPEALLVGKASQREGRLQQGNMRKNVRVLSRTSKFRRLRQLLRRRKKRWQGRRGGSRLEEEEEEEEEEEEEEEEEEEEEEEEEEEEEEEEEEEEEEEEEEEEEEEEE E

The code strings for all 20 phase differences are given in Column 3 in Table 5. Notice the reduction in the variation going from one phase to the next when a header and trailer are used.

**Table 5.**
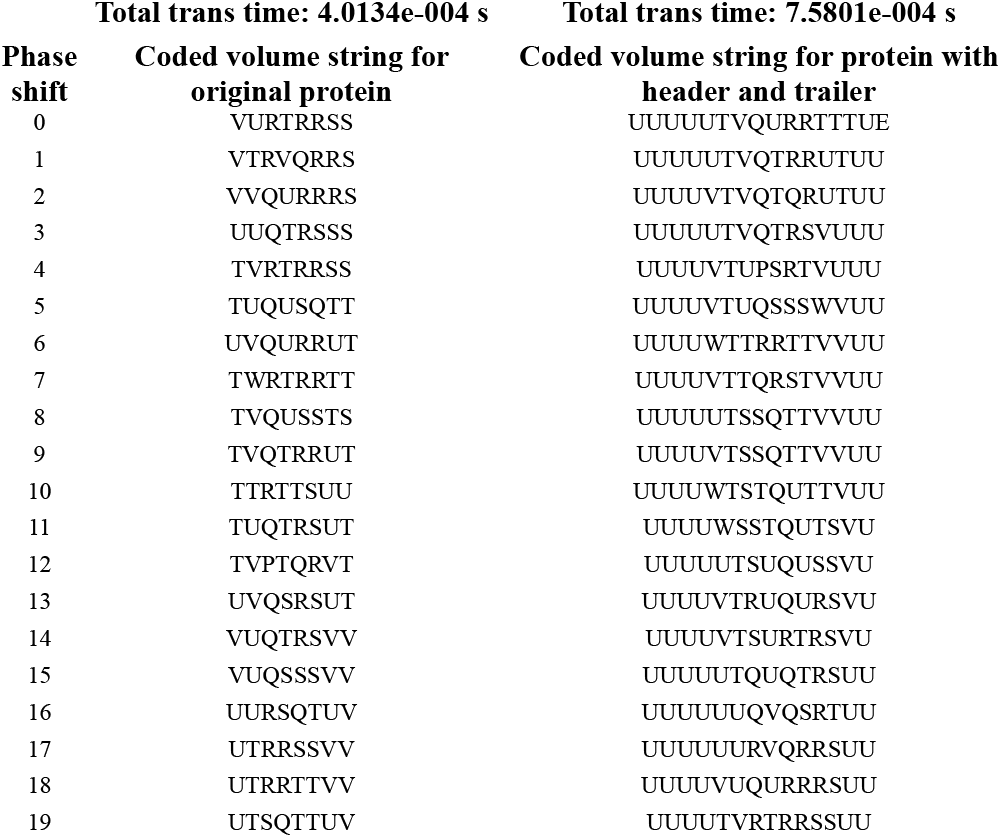
Coded volume strings for protein 5 in human proteome for phase difference of 0 to 19 residues with and without header and trailer

## 4. Discussion

1. One of the persistent issues in nanopore analysis of polymers is the speed with which an analyte translocates through a nanopore with a typical length of 5-10 nm. A large number of studies have focused on this, with a variety of methods proposed in theory and practice. In the present context the slowdown achieved should not be excessive as this would affect the total time required to process the large numbers of molecules in a sample. As noted earlier, samples from single cells can have as many as a billion molecules. In extreme cases, a single protein may have as many as 10^10^ copies while another may occur in a single copy. As the order in which the proteins enter the analyzing device, be it LC-MS or a nanopore, cannot be predicted the total analysis time becomes an important consideration. In [29] it is shown that a combination of opposing voltage and low or high pH can be used to slow down single bases, single AAs, short oligonucleotides, and short oligopeptides. This is possible because in all four cases a low or high pH value ensures that the mobility value will always have a sign that leads to a positive alpha in Equation 8c. Such is not the case when whole protein translocation is considered because inside the pore only K, R, Y, C, D, E, and H carry a charge, the other 13 AAs are internal residues (except when they are the first or last residue) and therefore do not carry a charge. Since K, R, and Y are positive and C, D, E, and H are negative and any of them can occur in a subsequence inside the pore there is a high degree of variation. In the present case a low pH value of 1 or 2 has been used because at this value D has a low negative mobility that is enough to slow down the pore sequence but not by so much as to delay it excessively. This is also the reason why a high value of pH has not been used. Thus with high pH slowdown will have to rely on R, which has a C_mult_ of 0.0239 at pH=13. This will result in excessive slowdown and cause total translocation times for the proteome to be several hours. Compare this with the C_mult_ value of 0.0022 for D at pH=1, which ensures that total translocation times for whole proteins in the proteome do not exceed ∼1 ms (Figure 5). In [26] unidirectional translocation of proteins is observed when the buffer includes guanidium chloride, which leads to approimately equal translocation times for all 20 AAs. However this also results in a loss of information about variations across the AAs, information that could be used to distinguish AAs or peptides from one another. With this method the average translocation time for a reside is 10 μs. With 1000 pores and an average of 400 residues per protein 10^9^ proteins can be counted in about 1.11 hrs. With biological pores the use of low or high pH solution for slowdown is predicated on the absence of any adverse effects on the pore protein.
2. Before a protein can enter the pore to translocate through it single file it has to be unfolded. Denaturation of proteins results in breaking of Cys-Cys bonds as well as unfolding of secondary structure. It is usually done by heating the protein or by using high pH values. In the present case heating is not an option.
3. Since a protein can enter the pore C-end or N-end first the direction of entry is of interest. In [26] a homopeptide with 10 D’s is used to determine the directionality of translocation. This is similar to the use of a header and trailer in Section 3. Here additionally the use of a header and trailer allows measuring the phase of the blockade signal with reference to the start of the protein. The resulting volume code string has a prefix and a suffix with the same code character. If a header and a trailer of different length and different AA are used the lengths of the prefix and suffix will be different (see Table 5).
4. Electroosmosis (EO), which causes a counterforce opposing the electrophoretic force [31], can sometimes be enough to reverse the direction of translocation. EO effects can be eliminated or greatly reduced by using a sheath of SDS [32], which effectively neutralizes the charge carried by residues on the wall of a biological pore. It can also be nullified by using a suitable buffer, for example see [19]. For these reasons EO effects have not been included in the model given here.
5. An analyte has to enter the pore within a reasonable amount of time. In [33] a nanopipette-based solution is given; the same effect can be approximated by a tapered *cis* chamber. Alternatively hydraulic pressure [34,35] can be used. However there is a trade-off: hydraulic pressure also prevents the flow of ions from *trans* to *cis*, thereby effectively halving the pore current; see [29].

## Supporting information

Supplementary information file includes following material, all pertaining to human proteome with Uniprot id UP000005640_9606: 1) List of 20598 protein sequences with id and length information; 2) List of volume code strings; 3) List of ids of length 6 in volume code strings for all proteins; 4) Table of translocation times and pore sequence volumes for protein number 5, id A6NL46.

